# Resolution of the circulating pathogenic phenotype of alpha-1 antitrypsin deficiency following liver transplantation

**DOI:** 10.64898/2025.11.29.691315

**Authors:** Nina Heyer-Chauhan, Riccardo Ronzoni, Ibrahim Aldobiyan, Elena Miranda, Aileen Marshall, John R. Hurst, James A. Irving, David A. Lomas

**Author notes:** Correspondence: N. Heyer-Chauhan. UCL Respiratory, Division of Medicine, University College London, Rayne Building, 5 University Street, London. WC1E 6JF, UK.

## Abstract

Alpha-1 antitrypsin is a member of the serpin family of protease inhibitors and normally found in monomeric form at high concentration in the circulation. Certain pathogenic variants of this protein self-assemble into flexible chains (polymers) that undergo toxic accumulation in the liver, underlying the development of liver disease, and a corresponding functional deficiency in circulation results in a susceptibility to chronic damage to the lungs and thereby COPD. A proportion of these polymers are detectable in the circulation. This study aimed to provide data on the dynamics of AAT monomers and polymers in the circulation after correction of a deficiency state by liver transplantation. The evolution of the circulating AAT phenotype was characterised quantitatively over time by analysis of blood samples taken pre- and post-liver transplant from three PiZZ AAT deficiency patients, using a panel of conformer-specific monoclonal antibodies and a novel ELISA with enhanced sensitivity for AAT polymers. Circulating wild-type M AAT was found to increase with a half-time of 29-39 h, attaining a clinically-defined putative ‘protective’ threshold level within 24-50 h post-transplant. Baseline circulating polymer levels ranged from 5-35 μg/mL with a half-time clearance of 3-12 h following transplant, and levels were indistinguishable from reference wild-type plasma 22 months post-surgery. The glycosylation profile and disappearance of detectable circulating polymers of AAT were consistent with their origin in, and canonical secretion by, the liver. To our knowledge this is the first post-transplant evaluation of the hybrid conformational profile of endogenous and donor tissue-derived AAT.

## Introduction

Alpha-1 antitrypsin (AAT), a member of the serpin family, is typically present in the human circulation at concentrations of 20-40 µM and is an inhibitor of proteases released by activated neutrophils. It is also associated with a conformational pathology known as alpha-1 antitrypsin deficiency (AATD). Around 95% of clinical cases of severe AATD are caused by the ‘Z’ variant of AAT (Glu342Lys) [1]. Homozygous individuals with a PiZZ genotype have plasma AAT levels that are 10-15% of the normal range, as most of the mutant protein is degraded or retained as chains of molecules, AAT polymers, within inclusions in the liver [2]. Physiologically, wild-type (or M) AAT protects lung tissue against proteolytic damage during an inflammatory response and low circulating levels are associated with early-onset panlobular emphysema. Conversely, polymer retention within the liver, where the protein is synthesised, is associated with neonatal hepatitis and cirrhosis [3] and adult liver disease [1]. Some polymers are secreted and can be detected in the plasma [4,5] where they are a biomarker of PiZZ individuals who will develop progressive liver disease [6–11] (Figure 1A). Formation of polymers in the liver involves an aberrant intermolecular interaction between AAT subunits in which a 4 kDa C-terminal region fails to self-incorporate and is instead donated to an adjacent molecule [12]. This process results in a conformational transition involving expansion of a central β-sheet, and mirrors a similar transition seen during inhibition of a target protease [13]. The different conformational states achieved by the protein, reflecting distinct molecular processes, can be probed by conformationally-selective monoclonal antibodies [12–16]. The AATD diagnosis pipeline typically involves an initial antibody-based quantification of circulating levels without distinguishing between conformers. While its use as a stratification tool may be controversial [17], a ‘protective threshold’ total AAT concentration of ∼11 µM in the circulation is generally considered to discriminate severe from mild deficiency and is a clinical target in augmentation therapy for PiZZ individuals [18,19]. The role of AAT as a protease inhibitor has been extensively characterised; anti-inflammatory roles have also been ascribed to AAT, such as opposing neutrophil chemotactic behaviour [20]. Conversely, AAT polymers are both non-functional as protease inhibitors and have been reported to display chemotactic activity [21–23].

**Figure 1:**
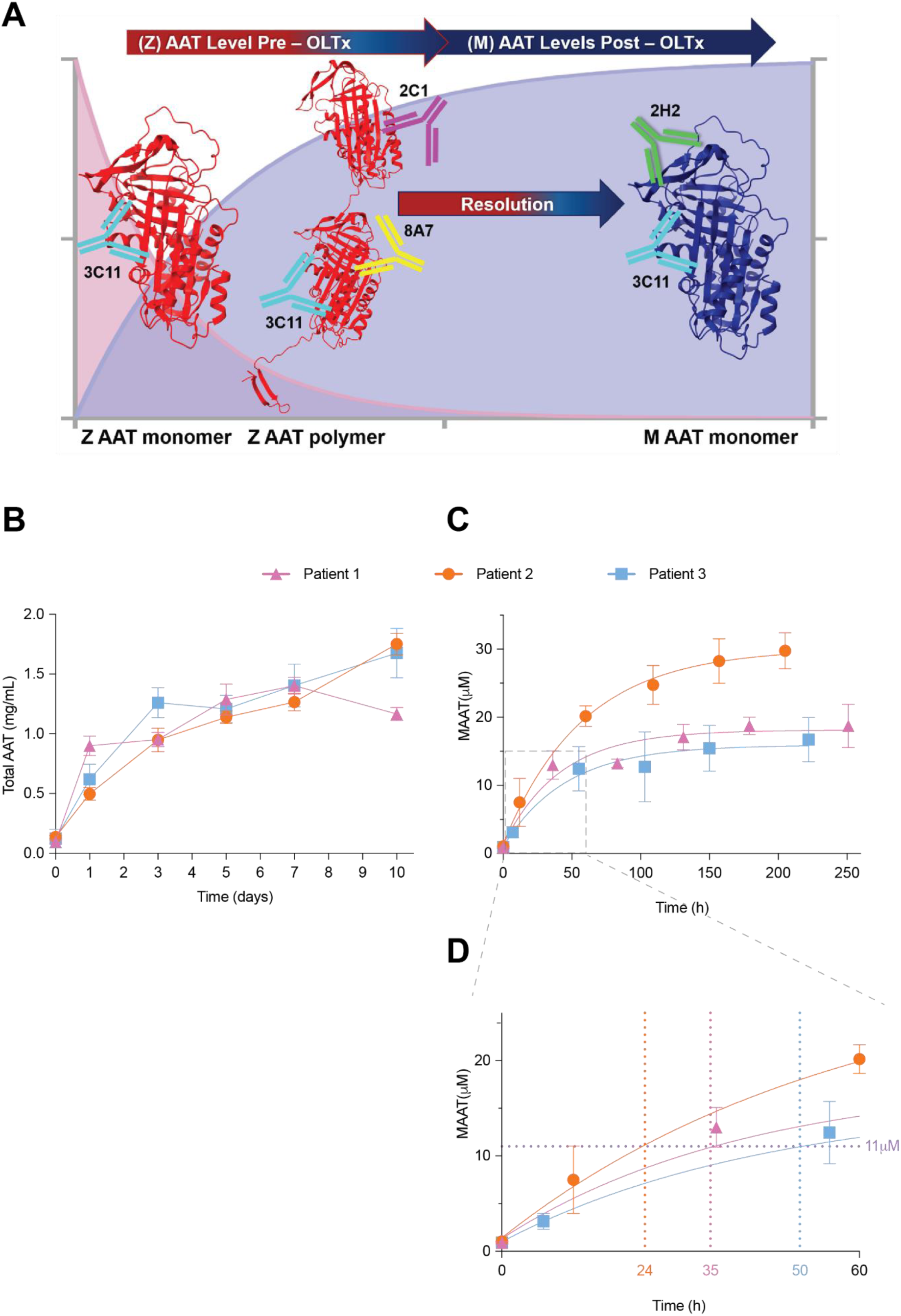
Increase in M AAT levels in plasma following liver transplantation. (A) Schematic representation of the specificities of the mAbs used in this study for M (blue) and Z (red) AAT conformations. (B) Total AAT for three individuals determined by sandwich ELISA using mAb 3C11 for detection (n=6). (C) M AAT for three individuals measured by sandwich ELISA using mAb 2H2 for detection (n=6). (D) Time taken (h) to reach the ‘protective threshold’ (11 µM, indicated by a lilac horizontal dotted line) for all three individuals. Values and error bars denote mean+/-SD.

The only approved therapy for end-stage lung and liver disease for individuals with AATD is transplantation, although several phase I/II clinical trials are underway evaluating novel strategies to supress or ablate the accumulation of intrahepatic inclusions of AAT or increase the level of circulating functional AAT produced by the liver [24]. However, there is little data quantitatively characterising the conformational AAT repertoire upon resolution of the deficiency state. The plasma of patients who have undergone orthotopic liver transplantation (OLTx), where the circulation will contain a combination of recipient-derived monomeric and polymeric Z AAT and nascent donor tissue-derived M AAT, provide an opportunity to obtain such information. Past studies have used isoelectric focussing to qualitatively characterise the exchange of circulating deficiency and non-deficiency variants upon OLTx by banding patterns in electrophoresis [25,26]. We have previously reported the quantitative decrease in circulating polymers in a single patient in the days following OLTx using a polymer-selective monoclonal antibody (mAb) 2C1 [4], and a more recent study by Kappe and coworkers expanded this evaluation with 5 PiZZ OLTx patients using the anti-Z AAT mAb LG96 [7]. However, the former study involved circulating polymer quantifications that were subject to intraoperative haemofiltration and the administration of blood products, preventing the acquisition of specific M AAT quantifications at that time, as they were subsumed within total AAT measurements. The latter study reported only one circulating polymer measurement following transplantation for each patient, performed at different times for each one (1-52 weeks post-OLTx), rather than monitoring the change in AAT conformers immediately following OLTx over time.

We have previously described several anti-AAT mAbs, including those that: recognise non-Z variants, including wild-type M AAT (mAb 2H2); distinguish monomer from polymer (mAbs 2C1 and 8A7); and recognise all conformations of M and Z AAT variants (mAb 3C11) [14–16,27]. Here, we have used these antibodies to determine the change in M and Z conformers of AAT following OLTx in three individuals with Z AATD. This has allowed us to determine the rates for the clearance of Z AAT polymers and the appearance of protective wild-type M AAT, characterise the biochemical properties of circulating Z AAT polymers, and confirm our previous studies showing that the liver is the source of detectable circulating polymers of AAT [5]. These findings offer an insight into the mechanisms involved in AATD-associated diseases and provide support for investigating the role of circulating polymer levels as a biomarker in patients with AATD.

## Methods

### Patients

The three subjects of this study had undergone OLTx. Patient-1 was a 66-year-old male with PiZ AATD and a 12-month history of portal hypertension, hepatic encephalopathy, gastro-oesophageal varices, ascites and hepatopulmonary syndrome. He had mild impairment of lung function (FEV_1_ 2.7L, 78% predicted; FVC 4.8L, 113% predicted) before his admission for OLTx. Patient-2 was a 42-year-old male with PiZZ AATD and non-alcoholic fatty liver disease (NAFLD). At the time of transplantation, he had hepatic encephalopathy, centripetal obesity and a history of calculous cholecystitis and anti-phospholipid syndrome. Patient-3 was a 48-year-old male who was diagnosed with PiZZ AATD in childhood and was asymptomatic until the development of ascites aged 44. All participants were enrolled in the London Alpha-1 Antitrypsin Deficiency Cohort Study and provided written informed consent. The study was conducted with ethics board approval (REC reference 13/LO/1085).

### Sample preparation

Plasma and explant liver tissue samples were collected from the three subjects. Z AAT polymers were extracted from liver tissue and isolated by immunoprecipitation [28]. Blood samples were collected immediately before, and on alternate days starting from, the day of transplantation (days 1-10) in 3.2% w/v citrate, and plasma was isolated by centrifugation at 1200 g for 30 minutes at 4°C. All samples were stored at -80°C until analysis. A long-term follow-up analysis was performed on day 669 post-OLTx in Patient-1.

### Antibodies and ELISA analysis

Quantification of AAT conformers was performed by sandwich ELISA following the general protocol described previously [29]. Polymers were detected by our standard assay using mAb 2C1 capture and mAb 3C11-HRP detection [6], or by a novel higher-sensitivity ELISA using mAb 2C1 capture and mAb 8A7-HRP detection, using our recently described 8A7 mAb [27]. The reference antigen used was a polymeric standard prepared by heating M AAT. The background signal from circulating M AAT at equivalent plasma dilution was subtracted prior to quantification. The limit of detection was 2.25 +/- 0.35 ng/ml and 0.73 +/- 0.16 ng/ml for the standard and high sensitivity assays respectively. M AAT was detected by mAb 3C11 capture and mAb 2H2 detection, and all AAT conformers (total AAT) by rabbit anti-human AAT polyclonal antibody (DAKO, Agilent Technologies, Inc. CA, USA) and mAb 3C11 detection [14–16]. The reference antigen used was a monomeric standard (M AAT) purified from PiMM AAT plasma. Quantified AAT polymer values were fitted using a non-linear least squares regression to a one-phase decay model and quantified M AAT values to a one phase association model. Statistical analyses were performed using Prism (Graphpad, Inc.).

### Extraction of liver AAT and analysis of N-linked glycosylation

Explant liver tissue was incubated in resuspension buffer [PBS supplemented with 1% v/v NP-40, Complete protease inhibitor cocktail (Roche Ltd., Hertfordshire, UK), 0.02% v/v sodium azide] and polymers were released by sonication at 4°C (Q500, QSonica, Newton, USA). Digestion with Peptide N-Glycosidase F (PNGase F, New England BioLabs Ltd, Hertfordshire, UK) or endoglycosidase H (Endo H, New England BioLabs Ltd, Hertfordshire, UK) was performed as described previously [28], following immunoprecipitation of polymers from samples with the 2C1 mAb, resolved by SDS-PAGE and visualised by western blot analysis with an anti-human AAT rabbit polyclonal antibody (DAKO) [5].

## Results

Plasma samples were provided by three PiZZ patients diagnosed with AATD who underwent OLTx, obtained on the day of surgery (day 0) and at days 1, 3, 5, 7 and 10 post-transplant. These samples were analysed by different sandwich ELISA assays to determine total, polymeric and M (wild-type) AAT in the circulation. By ten days post-surgery, plasma levels of total AAT had increased by approximately 10-fold from pre-transplantation levels to values within the normal range (1.5-3.5 mg/mL [30]) (Figure 1B and Table 1). A mAb selective for M AAT within the samples, mAb 2H2 [16], demonstrated that the rise in total AAT was due to an increase in the wild-type M protein (Figure 1C and Table 1). The estimated half-time for the increase in M AAT for subjects 1 to 3 following OLTx was 28.8 h +/- 6.6 h, 38.6 h +/- 7.1 h and 28.7 h +/- 10.4 h (estimate +/- standard error of the fit), respectively. Strikingly, by day 3 post OLTx all three individuals had M AAT levels that had exceeded the 11 µM putative protective threshold thought to confer lowered risk of COPD in AATD and generally used as a therapeutic target in replacement therapy [18,31,32] (Figure 1C).

**Table 1:**
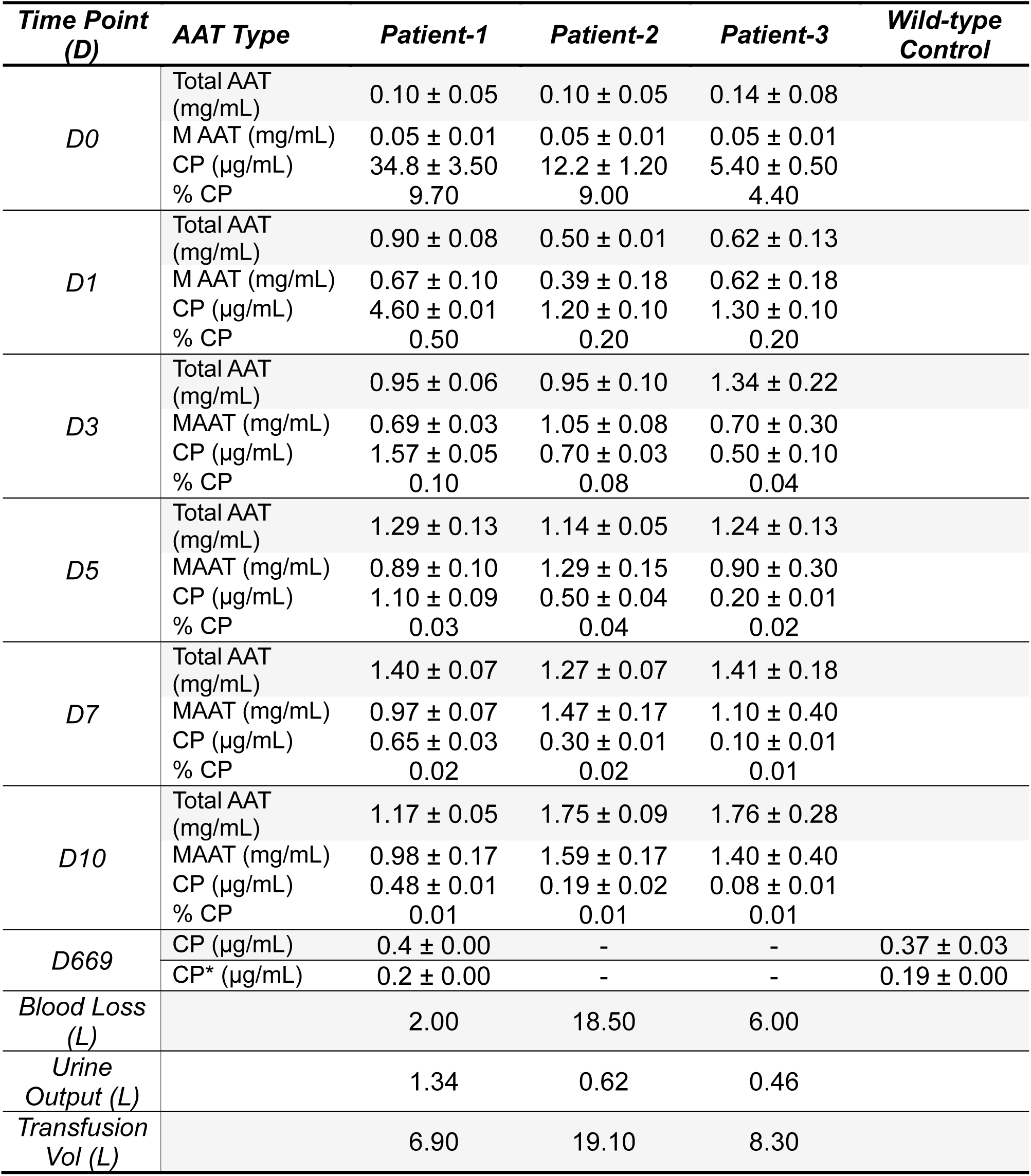
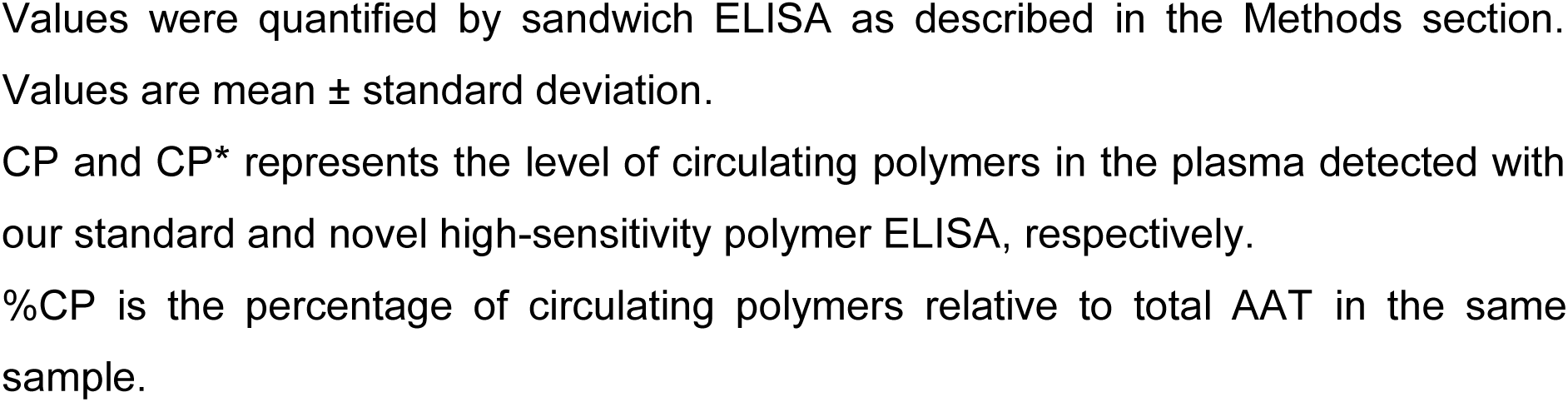
Plasma levels of AAT conformers pre- and post-OLTx and intraoperative fluid losses/gains.

| **Time Point (D)** | **AAT Type** | **Patient-1** | **Patient-2** | **Patient-3** | **Wild-type Control** |
| --- | --- | --- | --- | --- | --- |
| *D0* | Total AAT (mg/mL) | 0.10 ± 0.05 | 0.10 ± 0.05 | 0.14 ± 0.08 |  |
|  | M AAT (mg/mL) | 0.05 ± 0.01 | 0.05 ± 0.01 | 0.05 ± 0.01 |  |
|  | CP (µg/mL) | 34.8 ± 3.50 | 12.2 ± 1.20 | 5.40 ± 0.50 |  |
|  | % CP | 9.70 | 9.00 | 4.40 |  |
| *D1* | Total AAT (mg/mL) | 0.90 ± 0.08 | 0.50 ± 0.01 | 0.62 ± 0.13 |  |
|  | M AAT (mg/mL) | 0.67 ± 0.10 | 0.39 ± 0.18 | 0.62 ± 0.18 |  |
|  | CP (µg/mL) | 4.60 ± 0.01 | 1.20 ± 0.10 | 1.30 ± 0.10 |  |
|  | % CP | 0.50 | 0.20 | 0.20 |  |
| *D3* | Total AAT (mg/mL) | 0.95 ± 0.06 | 0.95 ± 0.10 | 1.34 ± 0.22 |  |
|  | MAAT (mg/mL) | 0.69 ± 0.03 | 1.05 ± 0.08 | 0.70 ± 0.30 |  |
|  | CP (µg/mL) | 1.57 ± 0.05 | 0.70 ± 0.03 | 0.50 ± 0.10 |  |
|  | % CP | 0.10 | 0.08 | 0.04 |  |
| *D5* | Total AAT (mg/mL) | 1.29 ± 0.13 | 1.14 ± 0.05 | 1.24 ± 0.13 |  |
|  | MAAT (mg/mL) | 0.89 ± 0.10 | 1.29 ± 0.15 | 0.90 ± 0.30 |  |
|  | CP (µg/mL) | 1.10 ± 0.09 | 0.50 ± 0.04 | 0.20 ± 0.01 |  |
|  | % CP | 0.03 | 0.04 | 0.02 |  |
| *D7* | Total AAT (mg/mL) | 1.40 ± 0.07 | 1.27 ± 0.07 | 1.41 ± 0.18 |  |
|  | MAAT (mg/mL) | 0.97 ± 0.07 | 1.47 ± 0.17 | 1.10 ± 0.40 |  |
|  | CP (µg/mL) | 0.65 ± 0.03 | 0.30 ± 0.01 | 0.10 ± 0.01 |  |
|  | % CP | 0.02 | 0.02 | 0.01 |  |
| *D10* | Total AAT (mg/mL) | 1.17 ± 0.05 | 1.75 ± 0.09 | 1.76 ± 0.28 |  |
|  | MAAT (mg/mL) | 0.98 ± 0.17 | 1.59 ± 0.17 | 1.40 ± 0.40 |  |
|  | CP (µg/mL) | 0.48 ± 0.01 | 0.19 ± 0.02 | 0.08 ± 0.01 |  |
|  | % CP | 0.01 | 0.01 | 0.01 |  |
| *D669* | CP (µg/mL) | 0.4 ± 0.00 | - | - | 0.37 ± 0.03 |
|  | CP* (µg/mL) | 0.2 ± 0.00 | - | - | 0.19 ± 0.00 |
| *Blood Loss (L)* |  | 2.00 | 18.50 | 6.00 |  |
| *Urine Output (L)* |  | 1.34 | 0.62 | 0.46 |  |
| *Transfusion Vol (L)* |  | 6.90 | 19.10 | 8.30 |  |

Values were quantified by sandwich ELISA as described in the Methods section. Values are mean ± standard deviation.
CP and CP\* represents the level of circulating polymers in the plasma detected with our standard and novel high-sensitivity polymer ELISA, respectively.
%CP is the percentage of circulating polymers relative to total AAT in the same sample.

Circulating polymers of AAT measured by ELISA using mAb 2C1 for capture and mAb 3C11-HRP for detection (lower limit of detection 2.25 +/- 0.35 ng/ml) [6,14] decreased as M AAT increased (Figure 2A and Table 1). This revealed an estimated half-life for the clearance of circulating polymers for the three subjects of 11.5 h +/- 1.1 h, 3.1 h +/- 0.4 h and 3.0 h +/- 0.2 h (estimate +/- standard error of the fit, n=6), respectively. These trends were also seen in the densitometric quantification of AAT isolated by immunoprecipitation of plasma samples from Patient-1, using the mAb 2C1 to isolate polymers and mAb 2H2 to isolate M AAT (Figure 2B and C), confirming the clearance of circulating polymers and the increase in M AAT over time.

**Figure 2:**
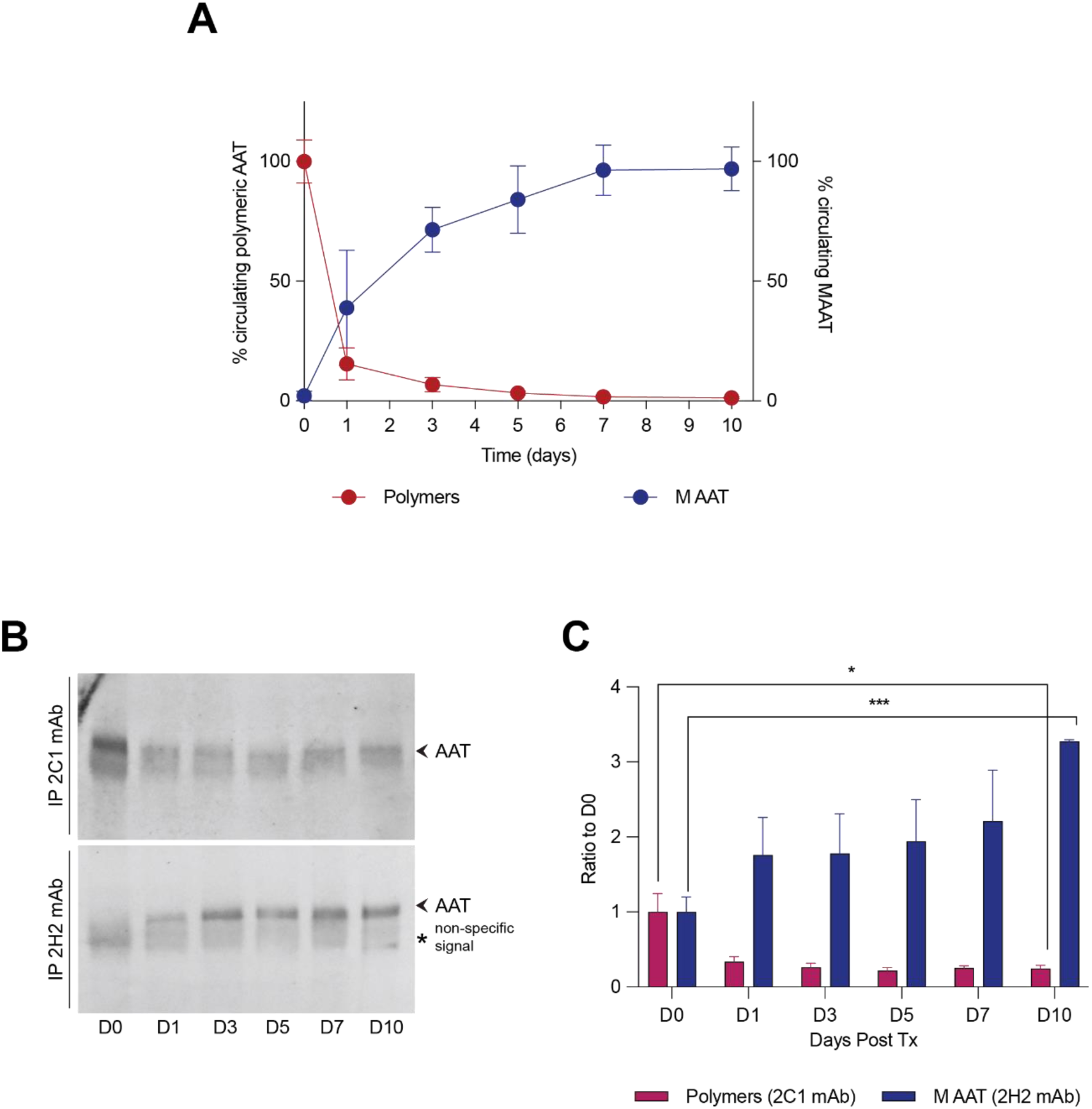
Change in circulating AAT conformers following liver transplantation. (A) Mean percentage of polymeric AAT (CP, circulating polymers) measured by ELISA with mAb 2C1 and M AAT by ELISA with mAb 2H2 for all three patients, relative to initial and final concentrations respectively (n=6). Values and error bars denote mean+/-SD. (B) Western blot of plasma samples immunoprecipitated with 2C1 to detect polymeric AAT or 2H2 mAb to detect M AAT. (C) Densitometric quantification of western blot in (B) (n=3, unpaired t test, *p<0.05, ****p <0.0005). Values and error bars denote mean+/-SD.

We have shown in the past that polymers of AAT are secreted from cells in culture models of AATD and that intracellular polymers show an immature, pre-Golgi N-glycosylation pattern while those in the extracellular millieu carry mature, post-Golgi N-glycans that are insensitive to Endo H digestion [5]. We applied the same analysis to the polymers immunoprecipitated from the plasma of Patient-1 using mAb 2C1, and we found them to be resistant to Endo H treatment, in keeping with a mature glycosylation profile and therefore reflecting exocytosis through the canonical secretory pathway. This contrasts with the immature, Endo H-sensitive high-mannose glycosylation nature of polymers extracted directly from a matched liver tissue sample (Figure 3). The present results agree with our previous findings, confirming the secretory origin of extracellular polymers and the endoplasmic reticulum as the organelle where intracellular polymers accumulate within liver cells.

**Figure 3:**
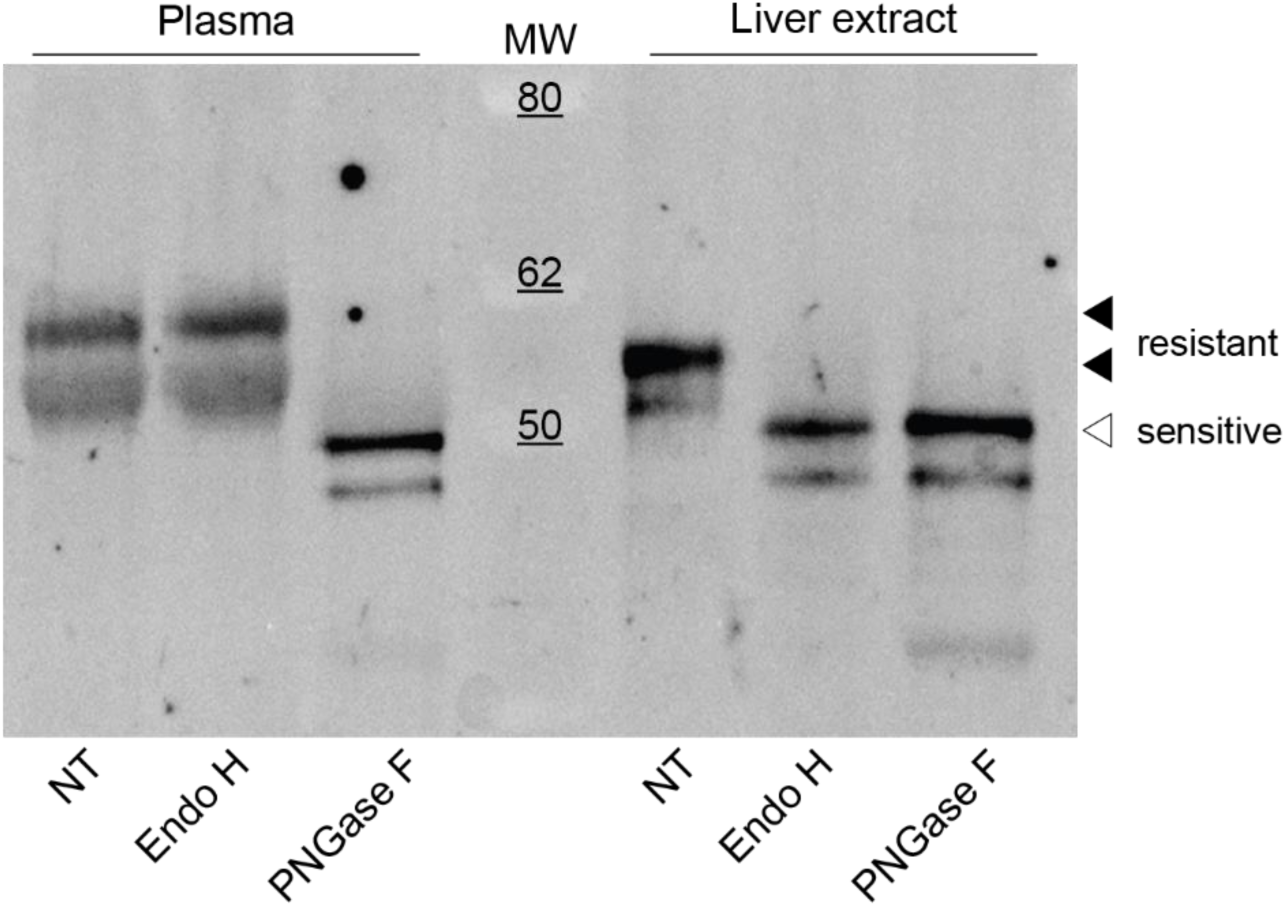
Glycosylation profiles of AAT from pre-OLTx plasma and explant liver tissue. Liver and plasma-derived AAT from Patient-1 were digested with Endo H (which cleaves glycans prior to maturation in the Golgi complex), PNGase F (which cleaves all glycan chains irrespective of maturity) or buffer control (NT). Black arrowheads point to AAT bands resistant to Endo H digestion, while white arrowheads indicate bands sensitive to Endo H.

Circulating polymers were still detectable at a low concentration in all three subjects 10 days following OLTx (Table 1). This raised the possibility that there may be an extra-hepatic source of circulating AAT polymers. An ELISA with enhanced sensitivity for AAT polymers with respect to the standard polymer detection assay was developed using a combination of the anti-polymer antibodies mAb 2C1 [14] for capture and mAb 8A7-HRP [27] for detection (lower limit of detection 0.73 +/- 0.16 ng/ml) (Figure 4A), and used to assess polymer levels 669 days post OLTx in Patient-1 (Figure 4B and Table 1). Both the standard and high-sensitivity assays provided a signal indistinguishable from the signal obtained with reference wild-type PiMM AAT plasma. This indicates that the liver is solely responsible for the production of the polymers detectable in the circulation.

**Figure 4:**
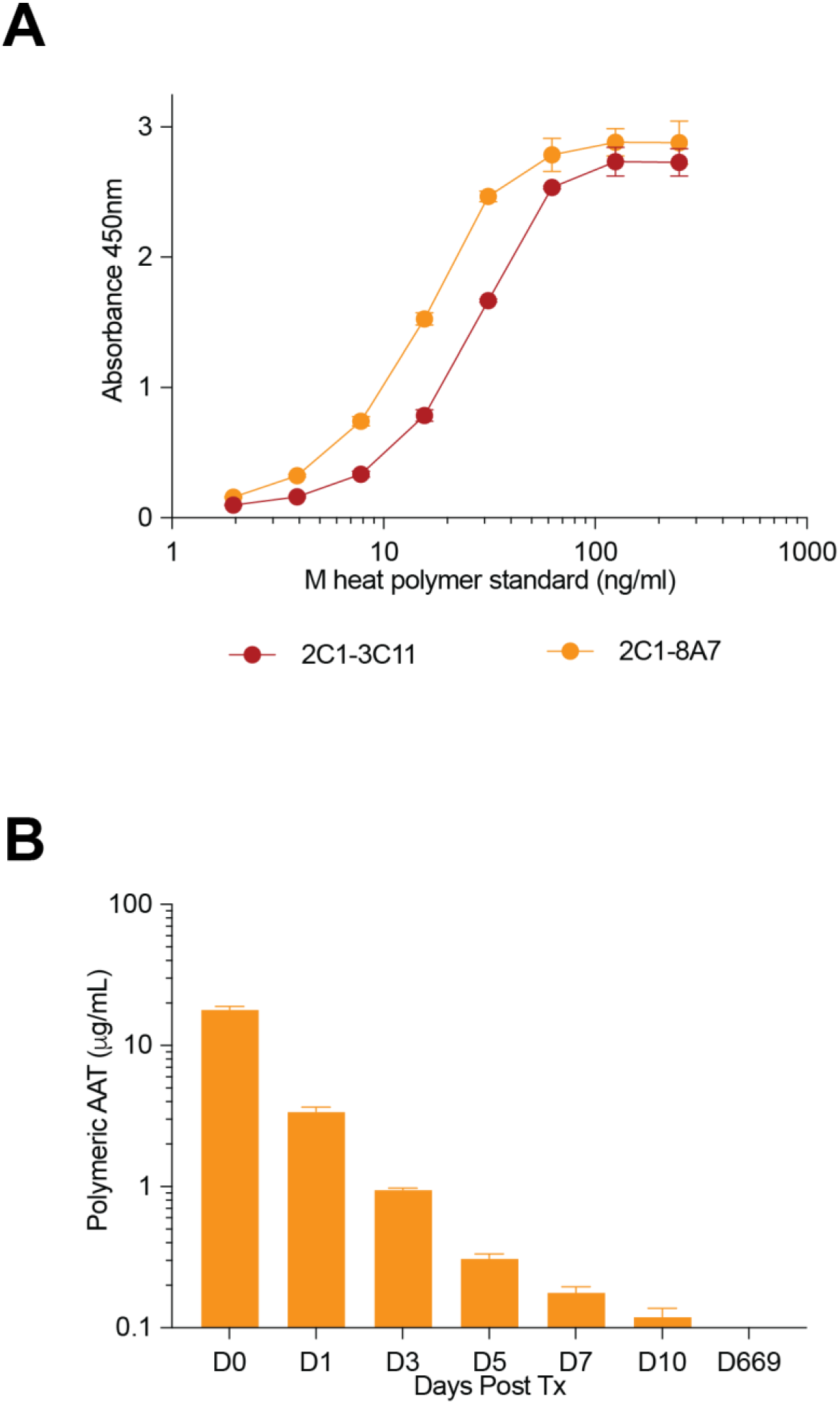
Polymeric AAT is absent from circulation months after liver transplantation. (A) Comparison of standard (2C1 capture, 3C11-HRP detection) and novel, high-sensitivity (2C1 capture, 8A7-HRP) sandwich ELISA assays for AAT polymer detection. The antigen used was a polymeric standard prepared by heating M AAT (n=3). (B) Plasma samples obtained from Patient 1 over time up to 669 days were evaluated with the high-sensitivity polymer ELISA. Values and error bars denote mean+/-SD.

## Discussion

AATD is a condition involving a protease–antiprotease imbalance in the lung, as well as an inflammatory disorder, brought about by insufficient levels of protective M AAT and accumulation of abnormal AAT predominantly in the liver, but also in the lung and other extrahepatic tissues [2]. Inflammation is predominantly neutrophil-driven, with several AAT–related mechanisms involved in potentiating this neutrophilic response [20]. Development of therapeutics for the treatment of AATD, targeting pathways involved in AATs anti-inflammatory effects and the strategies leading to prevention of intrahepatic polymer accumulation, all require an understanding of the dynamics of AAT production and clearance when considering endpoints.

The self-assembly of mutant AAT into polymer chains that condense as inclusion bodies within hepatocytes is one of the hallmarks of AATD. These polymers are the result of an aberrant self-assembly during production [2], are functionally inactive [33] and have pro-inflammatory properties [21,22,23]. They therefore constitute a pathologically important state of the protein. We have shown that the possession of a single PiZ allele results in detectable levels of polymers within the circulation [4,6] and that polymers are secreted by cells via the canonical secretory pathway [5,34]. On this basis we postulated that circulating AAT polymers may represent a surrogate of liver polymer load and therefore constitute a potential biomarker of disease [4,5]. Subsequent investigations have supported this hypothesis with circulating AAT polymers predicting individuals with adverse liver outcomes and in some cases correlating with more severe lung disease [6–11].

The method of quantification of circulating AAT employed in all these investigations was sandwich ELISA. However, it is critically important to note that the use of different capture and/or detection antibodies, as well as using different AAT reference protein standards can result in the calculation of different AAT values from the same plasma sample [14,15,35]. To further the field of AATD therapeutic development by ensuring commensurate comparison of data between different PiZZ cohort studies, we regard it as essential that a standardised form of ELISA screen for the quantification of circulating AAT polymers is agreed between different laboratories.

ELISA quantification of different AAT conformers involves different capture and detection antibody pairings. In this study, total AAT determination involved non-conformationally selective anti-AAT polyclonal for capture and mAb (3C11) for detection [15]. For M AAT determination, mAb 3C11 was used as the capture antibody and a monoclonal that does not bind to Z AAT, 2H2 [16], was used for detection. The epitope for the latter antibody localises to the intersection between β-sheet A and β-sheet C and binds this region preferentially for M AAT instead of the Z variant due to perturbation of this locus by the Z (Glu342Lys) substitution [16]. mAb 3C11, in contrast, recognises both the monomeric and polymeric forms of M and Z AAT with similar affinities [36].

AAT is predominantly expressed by hepatocytes with only low-level production by other cell types [37,38]. The hallmark of Z AAT production – polymers – were found to be undetectable in the plasma of a patient at a long follow-up time-point. Therefore, it can be presumed that the initial rise in circulating total AAT post-OLTx consists almost exclusively of wild-type M AAT with a minor contribution from resolving levels of monomeric and polymeric Z AAT in the days immediately following the transplant. This could account for small discrepancies seen between the quantified amounts of circulating total and M AAT in days 1-3 post-OLTx, where M AAT levels are on average 84% lower than the total AAT levels obtained from the same plasma sample (Table 1). However, it is probable that the different recognition capabilities of the capture and detection antibodies contribute to differences at later time points.

We have previously monitored the resolution of circulating Z AAT polymers for a single individual who had undergone OLTx, with intraoperative haemodialysis and substantial blood transfusion representing a caveat on the value obtained [4]. Here we have extended the existing polymer-detection assay with additional quantification of the nascent M AAT protein using the novel mAb 2H2, developed a novel ELISA with enhanced sensitivity using another novel mAb 8A7, and evaluated polymer clearance in three further individuals post-OLTx. All three individuals had levels of plasma AAT in the normal ‘healthy’ range ten days after hepatic transplantation and were all presenting a recognised pulmonary protective M AAT concentration by day three after surgery. Circulating polymers fell rapidly following OLTx. While the value in the first post-operative sample will be subject to the different blood losses and transfusion volumes that arose during transplantation, circulating polymers could still be detected, and were declining, at day 10, supporting the existence of systemic clearance mechanisms. This may be mediated by the LRP1 cell surface receptor, which is present on hepatocytes and monocytes [21,39,40]. However, it remains to be established whether this is the primary basis for the post-operative decline in polymers observed in our study. Our data provide evidence of a hepatic origin for the polymers of AAT that are detectable in the circulation, with the polymer signal around 22 months post-OLTx indistinguishable from that of the plasma of an PiMM individual. Moreover, the mature glycosylation profile shows that their presence in plasma is the result of active cellular secretion as opposed to cell rupture.

These data provide an understanding of the dynamics of M AAT and Z AAT in the circulation. They also demonstrate that the liver is exclusively responsible for the production of polymers detectable in the circulation and give mechanistic support for investigating the level of circulating polymers as a biomarker of AATD associated liver disease.

## Data availability

The data that support the findings of this study are provided in the paper and the raw data is available upon request from the corresponding author.

## Ethics approval and consent to participate

All research was conducted in accordance with both the Declarations of Helsinki and Istanbul. All research was conducted with ethics board approval (REC reference 13/LO/1085). All participants were enrolled in the London Alpha-1 Antitrypsin Deficiency Cohort Study and provided written informed consent.

## Authors’ contributions

NH-C, RR, AM, JRH, JAI and DAL designed the research; RR, NHC, EM, JAI and DAL analysed the data; NH-C, IA EM and RR performed the research; and NH-C, RR, EM, JAI and DAL wrote the paper; all authors edited and approved the paper.

## Consent for publication

All authors have consented to the publication of this manuscript.

## Permission to reproduce material from other sources

No material has been reproduced in this body of work.

## Clinical trial registration

This work is not part of a clinical trial.

## Acknowledgements

We thank the participants for providing samples used in this study. We would like to thank Reem Alluhibi, Shanaz Ahmad, Vivienne Hannon and The Liver Coordinator Team (especially Deidre Sexton and Michael McHugh) at Royal Free Hospital, Pond St, Hampstead, London NW3 2QG, U.K. for their assistance in the collection of the blood, tissue, clinical data and informed consents.

## Abbreviations

AAT: alpha-1 antitrypsin
AATD: alpha-1 antitrypsin deficiency
mAb: monoclonal antibody
Endo H: endoglycosidase H
M AAT: wild-type alpha-1 antitrypsin
OLTx: orthotopic liver transplantation
PiMM: individuals homozygous for wild-type M AAT
PiZZ: individuals homozygous for the Glu342Lys mutant of AAT
PNGase F: Peptide N-Glycosidase F
Z AAT: Glu342Lys mutant of AAT

## References

1. Strnad P, McElvaney NG, Lomas DA. Alpha1-Antitrypsin Deficiency. N Engl J Med. 2020;382:1443–55. 10.1056/NEJMra1910234

2. Lomas DA, Evans DL, Finch JT, Carrell RW. The mechanism of Z alpha 1-antitrypsin accumulation in the liver. Nature. 1992;357:605–7. 10.1038/357605a0

3. Cottrall K, Cook PJL, Mowat AP. Neonatal hepatitis syndrome and alpha-1-antitrypsin deficiency: an epidemiological study in South-East England. Postgrad Med J. 1974;50:376–80. 10.1136/pgmj.50.584.376

4. Tan L, Dickens JA, Demeo DL, Miranda E, Perez J, Rashid ST, et al. Circulating polymers in α1-antitrypsin deficiency. Eur Respir J. 2014;43:1501–4. 10.1183/09031936.00111213

5. Fra A, Cosmi F, Ordoñez A, Berardelli R, Perez J, Guadagno NA, et al. Polymers of Z α1-antitrypsin are secreted in cell models of disease. Eur Respir J. 2016;47:1005–9. 10.1183/13993003.00940-2015

6. Núñez A, Belmonte I, Miranda E, Barrecheguren M, Farago G, Loeb E, et al. Association between circulating alpha-1 antitrypsin polymers and lung and liver disease. Respir Res. 2021;22:244. 10.1186/s12931-021-01842-5

7. Sark AD, Fromme M, Olejnicka B, Welte T, Strnad P, Janciauskiene S, et al. The Relationship between Plasma Alpha-1-Antitrypsin Polymers and Lung or Liver Function in ZZ Alpha-1-Antitrypsin-Deficient Patients. Biomolecules. 2022;12:380. 10.3390/biom12030380

8. Teckman JH, Buchanan P, Blomenkamp KS, Heyer-Chauhan N, Burling K, Lomas DA. Biomarkers Associated With Future Severe Liver Disease in Children With Alpha-1-Antitrypsin Deficiency. Gastro Hep Adv. 2024;3:842–50. 10.1016/j.gastha.2024.04.010

9. Fromme M, Rademacher L, Amzou S, Cook CD, Zacharias I, Zhang L, et al. Association of circulating Z-polymer with adverse clinical outcomes and liver fibrosis in adults with alpha-1 antitrypsin deficiency. United European Gastroenterol J. 2024;12:1091–101. 10.1002/ueg2.12629

10. Suri A, Zhang Z, Neuschwander-Tetri B, Lomas DA, Heyer-Chauhan N, Burling K, et al. Fibrosis, biomarkers and liver biopsy in AAT deficiency and relation to liver Z protein polymer accumulation. Liver Int. 2024;44:3204–13. 10.1111/liv.16094

11. Kappe NN, Stolk J, Wout EFA van ’t, Janciauskiene SM, Hiemstra PS, van Hoek B. Validation of Biomarkers Z-AAT Polymers and Fibrinopeptide Aα-Val541 in Peripheral Blood of Patients With pi*ZZ Alpha-1 Antitrypsin Deficiency. Liver International Communications. 2025;6:e70022. 10.1002/lci2.70022

12. Aldobiyan I, Elliston ELK, Heyer-Chauhan N, Arold ST, Zhao L, Huntington B, et al. The mechanism of pathogenic α_1_ -antitrypsin aggregation in the human liver. Proc Natl Acad Sci USA. 2025;122:e2507535122. 10.1073/pnas.2507535122

13. Lowen SM, Waudby CA, Jagger AM, Aldobiyan I, Laffranchi M, Fra A, et al. High-resolution characterization of ex vivo AAT polymers by solution-state NMR spectroscopy. Science Advances. American Association for the Advancement of Science; 2025;11:eadu7064. 10.1126/sciadv.adu7064

14. Miranda E, Pérez J, Ekeowa UI, Hadzic N, Kalsheker N, Gooptu B, et al. A novel monoclonal antibody to characterize pathogenic polymers in liver disease associated with alpha1-antitrypsin deficiency. Hepatology. 2010;52:1078–88. 10.1002/hep.23760

15. Tan L, Perez J, Mela M, Miranda E, Burling KA, Rouhani FN, et al. Characterising the association of latency with α(1)-antitrypsin polymerisation using a novel monoclonal antibody. Int J Biochem Cell Biol. 2015;58:81–91. 10.1016/j.biocel.2014.11.005

16. Laffranchi M, Elliston EL, Miranda E, Perez J, Ronzoni R, Jagger AM, et al. Intrahepatic heteropolymerization of M and Z alpha-1-antitrypsin. JCI Insight. 2020;5:e135459, 135459. 10.1172/jci.insight.135459

17. Franciosi AN, Fraughen D, Carroll TP, McElvaney NG. Alpha-1 antitrypsin deficiency: clarifying the role of the putative protective threshold. Eur Respir J. 2022;59:2101410. 10.1183/13993003.01410-2021

18. Brantly ML, Wittes JT, Vogelmeier CF, Hubbard RC, Fells GA, Crystal RG. Use of a Highly Purified α1-Antitrypsin Standard to Establish Ranges for the Common Normal and Deficient α1-Antitrypsin Phenotypes. Chest. 1991;100:703–8. 10.1378/chest.100.3.703

19. Chapman KR, Burdon JGW, Piitulainen E, Sandhaus RA, Seersholm N, Stocks JM, et al. Intravenous augmentation treatment and lung density in severe α1 antitrypsin deficiency (RAPID): a randomised, double-blind, placebo-controlled trial. Lancet. 2015;386:360–8. 10.1016/S0140-6736(15)60860-1

20. McCarthy C, Reeves EP, McElvaney NG. The Role of Neutrophils in Alpha-1 Antitrypsin Deficiency. Ann Am Thorac Soc. 2016;13 Suppl 4:S297–304. 10.1513/AnnalsATS.201509-634KV

21. Parmar JS, Mahadeva R, Reed BJ, Farahi N, Cadwallader KA, Keogan MT, et al. Polymers of alpha(1)-antitrypsin are chemotactic for human neutrophils: a new paradigm for the pathogenesis of emphysema. Am J Respir Cell Mol Biol. 2002;26:723–30. 10.1165/ajrcmb.26.6.4739

22. Mulgrew AT, Taggart CC, Lawless MW, Greene CM, Brantly ML, O’Neill SJ, et al. Z alpha1-antitrypsin polymerizes in the lung and acts as a neutrophil chemoattractant. Chest. 2004;125:1952–7. 10.1378/chest.125.5.1952

23. Mahadeva R, Atkinson C, Li Z, Stewart S, Janciauskiene S, Kelley DG, et al. Polymers of Z alpha1-antitrypsin co-localize with neutrophils in emphysematous alveoli and are chemotactic in vivo. Am J Pathol. 2005;166:377–86. 10.1016/s0002-9440(10)62261-4

24. Alluhibi R, Marshall A, Lomas DA, Hurst JR. A cure for alpha-1? Novel therapeutics in alpha-1 antitrypsin deficiency. European Respiratory Journal [Internet]. European Respiratory Society; 2025 [cited 2025 Nov 17];66. 10.1183/13993003.01101-2025

25. Hood JM, Koep LJ, Peters RL, Schröter GP, Weil R, Redeker AG, et al. Liver transplantation for advanced liver disease with alpha-1-antitrypsin deficiency. N Engl J Med. 1980;302:272–5. 10.1056/NEJM198001313020505

26. van Furth R, Kramps JA, van der Putten AB, Krom RA, Gips CH. Change in alpha 1-antitrypsin phenotype after orthotopic liver transplant. Clin Exp Immunol. 1986;66:669–72.

27. Ronzoni R, Aldobyian IF, Miranda E, Heyer-Chauhan N, Elliston EL, Pérez J, et al. Susceptibility of alpha1 antitrypsin deficiency variants to polymer blocking therapy. JCI Insight. 2025;e194354. 10.1172/jci.insight.194354

28. Faull SV, Elliston ELK, Gooptu B, Jagger AM, Aldobiyan I, Redzej A, et al. The structural basis for Z α1-antitrypsin polymerization in the liver. Sci Adv. 2020;6:eabc1370. 10.1126/sciadv.abc1370

29. Belorgey D, Irving JA, Ekeowa UI, Freeke J, Roussel BD, Miranda E, et al. Characterisation of serpin polymers in vitro and in vivo. Methods. 2011;53:255–66. 10.1016/j.ymeth.2010.11.008

30. Miravitlles M, Herr C, Ferrarotti I, Jardi R, Rodriguez-Frias F, Luisetti M, et al. Laboratory testing of individuals with severe alpha1-antitrypsin deficiency in three European centres. Eur Respir J. 2010;35:960–8. 10.1183/09031936.00069709

31. Soy D, de la Roza C, Lara B, Esquinas C, Torres A, Miravitlles M. Alpha-1-antitrypsin deficiency: optimal therapeutic regimen based on population pharmacokinetics. Thorax. 2006;61:1059–64. 10.1136/thx.2005.057943

32. Franciosi AN, Hobbs BD, McElvaney OJ, Molloy K, Hersh C, Clarke L, et al. Clarifying the Risk of Lung Disease in SZ Alpha-1 Antitrypsin Deficiency. Am J Respir Crit Care Med. American Thoracic Society -AJRCCM; 2020;202:73–82. 10.1164/rccm.202002-0262OC

33. Lomas DA, Finch JT, Seyama K, Nukiwa T, Carrell RW. Alpha 1-antitrypsin Siiyama (Ser53-->Phe). Further evidence for intracellular loop-sheet polymerization. J Biol Chem. 1993;268:15333–5.

34. Ronzoni R, Heyer-Chauhan N, Fra A, Pearce AC, Rüdiger M, Miranda E, et al. The molecular species responsible for α1 -antitrypsin deficiency are suppressed by a small molecule chaperone. FEBS J. 2021;288:2222–37. 10.1111/febs.15597

35. Janciauskiene S, Welte T, Lehmann M. An Enzyme-Linked Immunosorbent Assay (ELISA) for Quantification of Circulating Pi*Z Alpha1-Antitrypsin Polymers. Methods Mol Biol. 2024;2750:113–22. 10.1007/978-1-0716-3605-3_11

36. Ordóñez A, Pérez J, Tan L, Dickens JA, Motamedi-Shad N, Irving JA, et al. A single-chain variable fragment intrabody prevents intracellular polymerization of Z α1-antitrypsin while allowing its antiproteinase activity. FASEB J. 2015;29:2667–78. 10.1096/fj.14-267351

37. Mornex JF, Chytil-Weir A, Martinet Y, Courtney M, LeCocq JP, Crystal RG. Expression of the alpha-1-antitrypsin gene in mononuclear phagocytes of normal and alpha-1-antitrypsin-deficient individuals. J Clin Invest. 1986;77:1952–61. 10.1172/JCI112524

38. van ‘t Wout EFA, Dickens JA, van Schadewijk A, Haq I, Kwok HF, Ordóñez A, et al. Increased ERK signalling promotes inflammatory signalling in primary airway epithelial cells expressing Z α1-antitrypsin. Hum Mol Genet. 2014;23:929–41. 10.1093/hmg/ddt487

39. Joslin G, Fallon RJ, Bullock J, Adams SP, Perlmutter DH. The SEC receptor recognizes a pentapeptide neodomain of alpha 1-antitrypsin-protease complexes. J Biol Chem. 1991;266:11282–8.

40. Strickland DK, Muratoglu SC, Antalis TM. Serpin-Enzyme Receptors LDL Receptor-Related Protein 1. Methods Enzymol. 2011;499:17–31. 10.1016/B978-0-12-386471-0.00002-X

