## Supplemental Fig 2B & Fig 3 for "Resolution of the circulating pathogenic phenotype of alpha-1 antitrypsin deficiency following liver transplantation"

### Uncropped blots for Figure 2B and Figure 3 shown in the main text

Uncropped blot for Figure 2B IP of plasma samples with 2C1 mAb and IP of plasma samples 2H2 mAb in the main text:

Loading Order:

| | Top Blot (2C1) | | Lower Blot (2H2) |
|---|---|---|---|
| 1. | IP 2C1 Day 0 | 7. | IP 2H2 Day 0 |
| 2. | IP 2C1 Day 1 | 8. | IP 2H2 Day 1 |
| 3. | IP 2C1 Day 3 | 9. | IP 2H2 Day 3 |
| 4. | IP 2C1 Day 5 | 10. | IP 2H2 Day 5 |
| 5. | IP 2C1 Day 7 | 11. | IP 2H2 Day 7 |
| 6. | IP 2C1 Day 10 | 12. | IP 2H2 Day 10 |

Uncropped blot showing IP of plasma samples with mAb 2C1 from the main text in Figure 2B

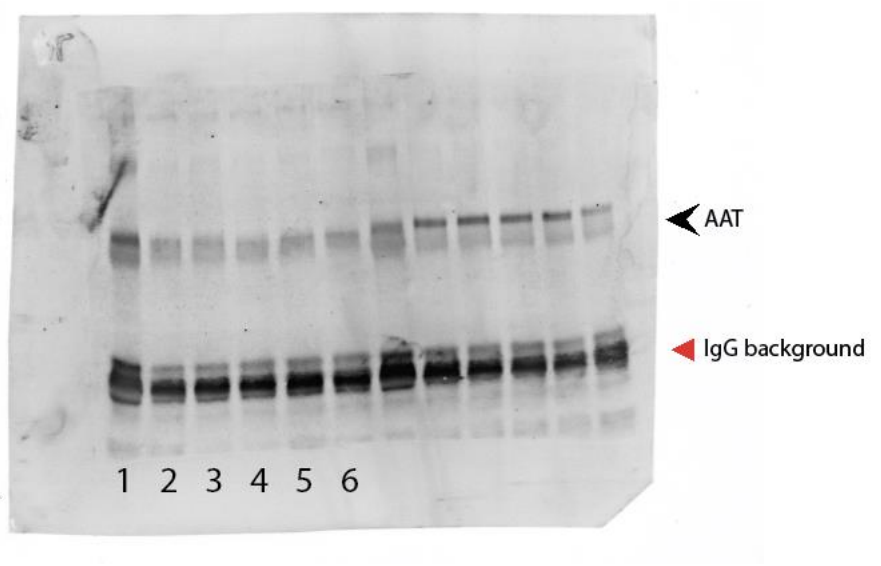


Uncropped blot showing IP of plasma samples with mAb 2H2 from the main text in Figure 2B

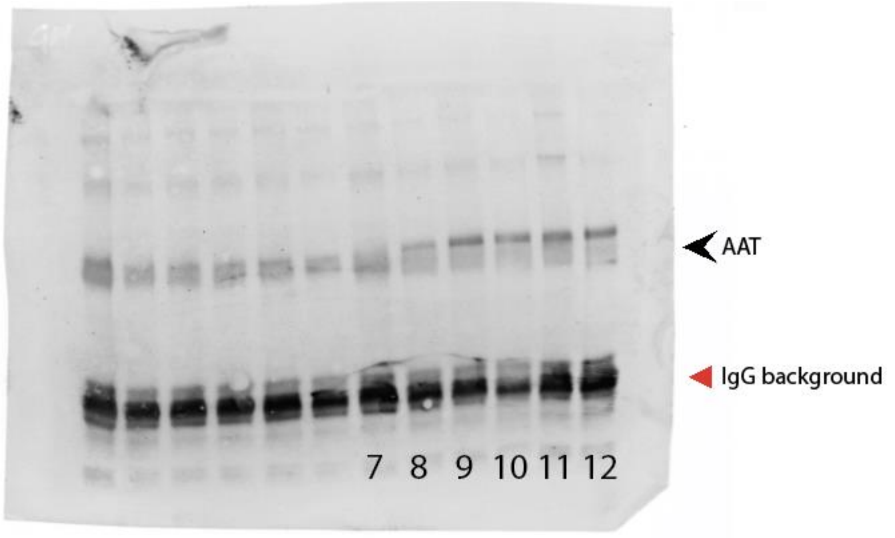

Uncropped Western Blot for Figure 3: “Glycosylation profiles of AAT from pre-OLTx plasma and explant liver tissue.” In the main text:

Loading Order:

1. Molecular Weight Marker
2. PiZZ plasma not treated
3. PiZZ plasma Endo H treated
4. PiZZ plasma PNGase F treated
5. Molecular Weight Marker
6. PiZZ liver extract not treated
7. PiZZ liver extract Endo H treated
8. PiZZ liver extract PNGase F treated
9. Molecular Weight Marker

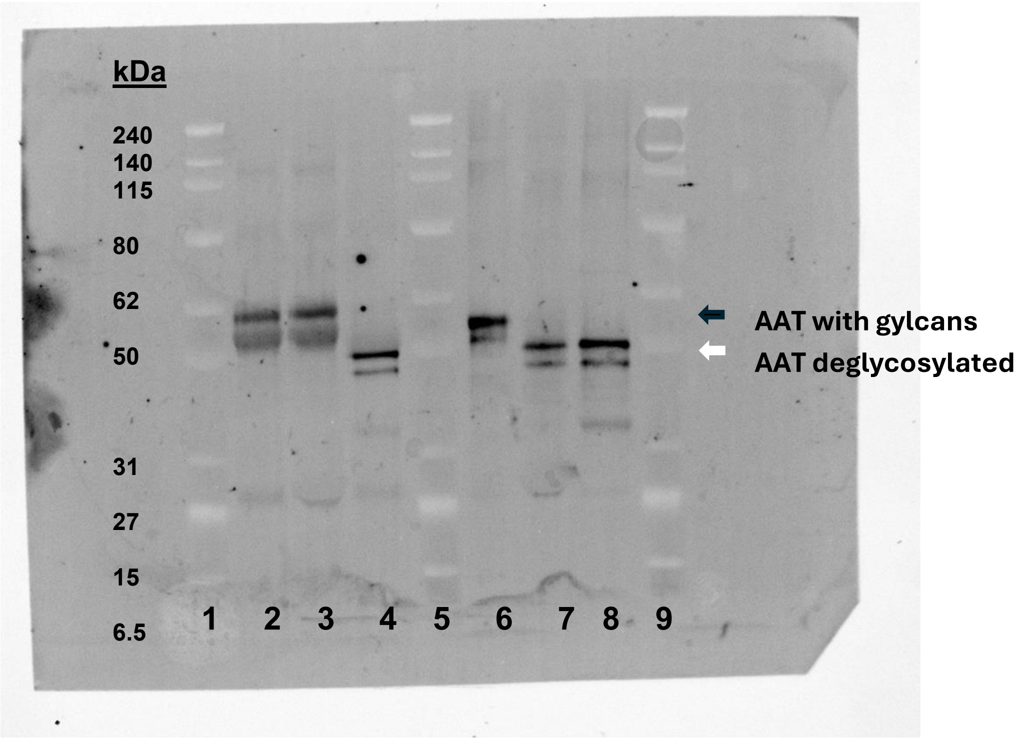
